# SF3A2 deacetylation at lysin 10 mitigates myocardial pathological remodeling by reprogramming cardiomyocyte metabolism

**DOI:** 10.64898/2025.12.26.696635

**Authors:** Renpeng Guo, Zepeng Zhang, He Zhang, Yue Yang, Ming Qiao, Jiawei Zhang, Yisa Wang, Jinjin Chen, Jing Li, Mingxia Wu, Kexin Yang, Yangyang Qing, Haisi Dong, Xiangyan Li, Daqing Zhao, Qingxia Huang

## Abstract

**Background:** Dysregulation of splicing factor-mediated pre-mRNA alternative splicing (AS) underlies the progression of complex diseases, but the specific involvement of AS in myocardial pathological remodeling remains unclear. This study aimed to elucidate the role of splicing factor 3A, subunit 2 (SF3A2) in the development of myocardial pathological remodeling by regulating AS and its effect on metabolic reprogramming.

**Methods:** We evaluated the function and mechanism of SF3A2 in myocardial pathological remodeling using murine models induced by myocardial infarction (MI) and isoproterenol. The gain– and loss-of-function models (AAV9-cTNT delivery-mediated knock-in and lentivirus-mediated knockdown) combined with acetylomics, metabolomics, lipidomics, ^13^C-glucose tracing and immunoprecipitation coupled with liquid chromatography-tandem mass spectrometry (IP-MS) analyses were used to demonstrate the mechanism of SIRT7-mediated SF3A2 deacetylation at lysin 10 (K10). Unbiased RNA sequencing and AS analysis were conducted to identify the downstream effector of SF3A2.

**Results:** SF3A2 acetylation at lysine 10 was upregulated in cardiomyocytes in response to myocardial pathological remodeling. Functionally, the knock-in of a deacetylation-mimetic SF3A2 (K10R) mutant attenuated myocardial hypertrophy, fibrosis, and improved heart function. These phenotypes were accompanied by a redirection of glucose flux from anaerobic glycolysis to the TCA cycle, along with a promotion of fatty acid β-oxidation by inhibiting CD36 translocation and decreasing ACLY activity. Mechanistically, SF3A2 mitigated metabolic reprogramming and mitochondrial dysfunction by regulating Nnt AS, a process involving the SF3A2-interacting protein Ddx1. IP-MS further identified SIRT7 as the deacetylase for SF3A2, which was found to be activated by ginsenoside Rg1 in a high-throughput screen, leading to improved heart function.

**Conclusions:** Overall, our findings establish the SIRT7-SF3A2-Nnt axis as a critical regulator in metabolic reprogramming to delay pathological myocardial remodeling. Promoting SF3A2 deacetylation, mediated by AAV9 delivery or pharmacological activation of SIRT7, improved heart function, suggesting its potential as a therapeutic target for pathological myocardial remodeling.

## Introduction

Myocardial infarction (MI) is a leading cause of death worldwide^1^. While advancements in percutaneous coronary intervention have reduced acute mortality, myocardial pathological remodeling remains a major obstacle to improve long-term survival^2, 3^. Myocardial pathological remodeling is a maladaptive structural and functional alterations in response to MI, characterized by hypertrophy, fibrosis, and heart failure^4^. Current first-line pharmacological therapies, such as renin-angiotensin-aldosterone system inhibitors and β-blockers, offer limited efficacy in reversing pathological remodeling and fail to reduce the incidence of heart failure^1, 5^. Therefore, it is important to advance our understanding of molecular mechanisms for preventing or reversing myocardial pathological remodeling.

Metabolic reprogramming in cardiomyocytes is a hallmark of myocardial pathological remodeling^6^. The onset of ischemia/reperfusion (I/R) after revascularization for MI is accompanied by a metabolic shift, driven by alterations in the peri-myocyte milieu (pressure overload, immune response and inflammation) and substrate availability, which collectively upregulate genes favoring anaerobic glycolysis over fatty acid oxidation^7–9^. During pathological remodeling, the translocation of CD36 to the plasma membrane enhances fatty acid uptake from the circulation^10^. However, concurrently impaired mitochondrial β-oxidation prevents the efficient utilization of these fatty acids, leading to their intracellular accumulation and subsequent lipotoxicity^11^. Mitochondrial dysfunction alters TCA flux, increasing the accumulation of citrate for ATP-citrate lyase (ACLY) to generate nuclear acetyl-CoA^12^. This acetyl-CoA surplus in nucleus serves as an acetyl donor and drives widespread protein hyperacetylation^13^. In addition to substrate-level changes, a NAD^+^ depletion impairs SIRT deacetylase function, creating an imbalance that favors acetyl-group accumulation, thereby altering the cellular acetylome and contributing to heart dysfunction^14^. Normalization of acetylation represents a crucial therapeutic direction for improving myocardial pathological remodeling.

Dysregulation of pre-mRNA alternative splicing (AS) underlies the development and progression of many diseases^15–17^. As a key splicing factor of the U2 snRNP complex, splicing factor 3A, subunit 2 (SF3A2) is recruited to pre-mRNA during AS, where it recognizes and binds the branch point sequence and the 3’ splice site^18^. This facilitates the assembly of a catalytically active spliceosome that carries out the transesterification reactions, thereby producing mature mRNA for protein translation^19^. Genome-wide association studies have revealed that SF3A2 is associated with both the onset and prognosis of myocardial infarction^20^. Additionally, the acetylation of SF3A2 in response to ischemic stress regulates the AS of mitochondrial genes^21, 22^. However, the function of SF3A2 in metabolic reprogramming and myocardial pathological remodeling and the underlying mechanisms remain poorly understood.

Here we report that acetylation modification at lysine 10 in SF3A2 is a potential driver event in myocardial pathological remodeling. SF3A2 deacetylation knock-in rats exhibited restored heart function and mitochondrial oxidative phosphorylation (OXPHOS). This phenotype concomitantly redirected glucose flux toward TCA cycle over anaerobic glycolysis and promoting fatty acid β-oxidation. Mechanistically, SIRT7, as a deacetylase, disassociates from SF3A2 in response to myocardial pathological remodeling. SF3A2 hyperacetylation enhances its interaction with Ddx1 to AS, which promotes the exon skipping of Nnt pre-mRNA and ultimately leads to metabolic reprogramming. Furthermore, pharmacological activator of SIRT7 with a lead compound suggests a potential therapeutic strategy against myocardial pathological remodeling. Our findings describe the SIRT7-SF3A2 acetylation axis as a novel pathway in regulating metabolic reprogramming to protect against myocardial pathological remodeling.

## Methods

### Data Availability

The data, analytical methods, and study materials supporting the findings of this study are available from the corresponding author upon reasonable request. The mass spectrometry proteomics and acetylomic data have been deposited to the ProteomeXchange Consortium (http://proteomecentral.proteomexchange.org) via the iProX partner repository with the dataset identifier PXD069931. The transcriptome sequencing data are available in the Gene Expression Omnibus (GEO; https://www.ncbi.nlm.nih.gov/geo/) under accession number GSE236472.

The detailed methods used in this study are provided in the Supplemental Material.

## Results

### 1 Identification of SF3A2 as a driving factor of myocardial pathological remodeling

To identify the profile of early acetylated modification underlying pathological remodeling, we conducted acetylomics in H9c2 cells. This model can be differentiated into a cell type which recapitulates many key features of cardiomyocytes. As shown in Fig. 1A, PCA analysis revealed distinct clustering among control, oxygen and glucose deprivation (OGD), and OGD/reperfusion (OGD/R) groups in the acetylomics. Venn diagram analysis identified 57 commonly up-regulated and 24 down-regulated acetylated proteins between the OGD *vs.* control and OGD/R *vs.* control comparisons (Fig. S1A). KEGG pathway and heatmap enrichment analysis of differentially acetylated proteins revealed that hyperacetylated proteins in both OGD and OGD/R groups were significantly enriched in pathways of longevity, spliceosome and ECM−receptor interaction, while deacetylated proteins were enriched in oocyte meiosis and cholesterol metabolism pathways (Fig. 1B-1C). We also conducted a protein-protein interaction (PPI) analysis, which revealed that spliceosome-associated proteins serve as central mediators among differentially acetylated proteins in cardiomyocytes following OGD and OGD/R incubation (Fig. 1D). The volcano plot results further confirmed that spliceosome-associated proteins (SF3A2, U2AF2 and CWC15) were among the most significantly acetylated proteins (Fig. 1E and Fig. S1B). Based on these results, we targeted spliceosome-associated proteins and validated the acetylomic results by co-immunoprecipitation (co-IP). Lysates of H9c2 cells and primary cardiomyocytes were immunoprecipitated with an anti-pan acetylation (Ac-K) antibody and probed with anti-U2AF2, –CWC15 or –SF3A2 antibody. Correspondingly, the primary cardiomyocyte lysates were also immunoprecipitated with an anti-SF3A2 antibody and probed with an anti-Ac-K antibody. The co-IP experiment showed that SF3A2 and Ac-K are complexed in cardiomyocytes after OGD or OGD/R incubation (Fig. 1F). To determine whether SF3A2 acetylation is associated with myocardial pathological remodeling, we established a rat model of myocardial ischemia followed by reperfusion for 4 hours, 1 day, 7 days or 14 days (Fig. 1G). Echocardiographic and hemodynamic assessments revealed a temporal evolution of heart dysfunction: systolic dysfunction at day 1, reduced aortic arch flow velocity at day 7, and diastolic dysfunction at day 14 post-reperfusion (Fig. 1H and Fig. S1C-S1D). The serum levels of myocardial injury markers, creatine kinase (CK) and lactate dehydrogenase (LDH), peaked at 4 hours of reperfusion (Fig. S1E). H&E, Masson and WGA staining showed a time-dependent aggravation of disturbed myofibrillar arrangement, myofibrillar swelling and myocardial fibrosis, accompanied by marked cardiomyocyte hypertrophy after 7 days of reperfusion (Fig. S2A). These results indicate an ongoing process of myocardial pathological remodeling during the 14-days I/R period. More importantly, SF3A2 acetylation exhibited a slight increase after 1 hour of ischemia, whereas a pronounced increase was observed following reperfusion, accompanied by a reduction in the Bcl-2/Bax ratio (Fig. 1I). Confocal microscopy also conformed the co-localization of SF3A2 and Ac-K in nucleus on post-reperfusion (Fig. 1J and Fig. S2B). Next, to determine whether SF3A2 hyperacetylation is conserved in pathological remodeling, we employed an *in vitro* model of adult mouse cardiomyocytes (ACMs) that more faithfully recapitulates aspects of cardiomyocyte biology. As shown in Fig. 1K, the SF3A2 hyperacetylation was also observed in ACMs isolated from a mouse model of isoproterenol (ISO)-induced cardiac hypertrophy. These results demonstrated that an increase in SF3A2 acetylation is a general phenotype underlying the myocardial pathological remodeling.

**Fig. 1.**
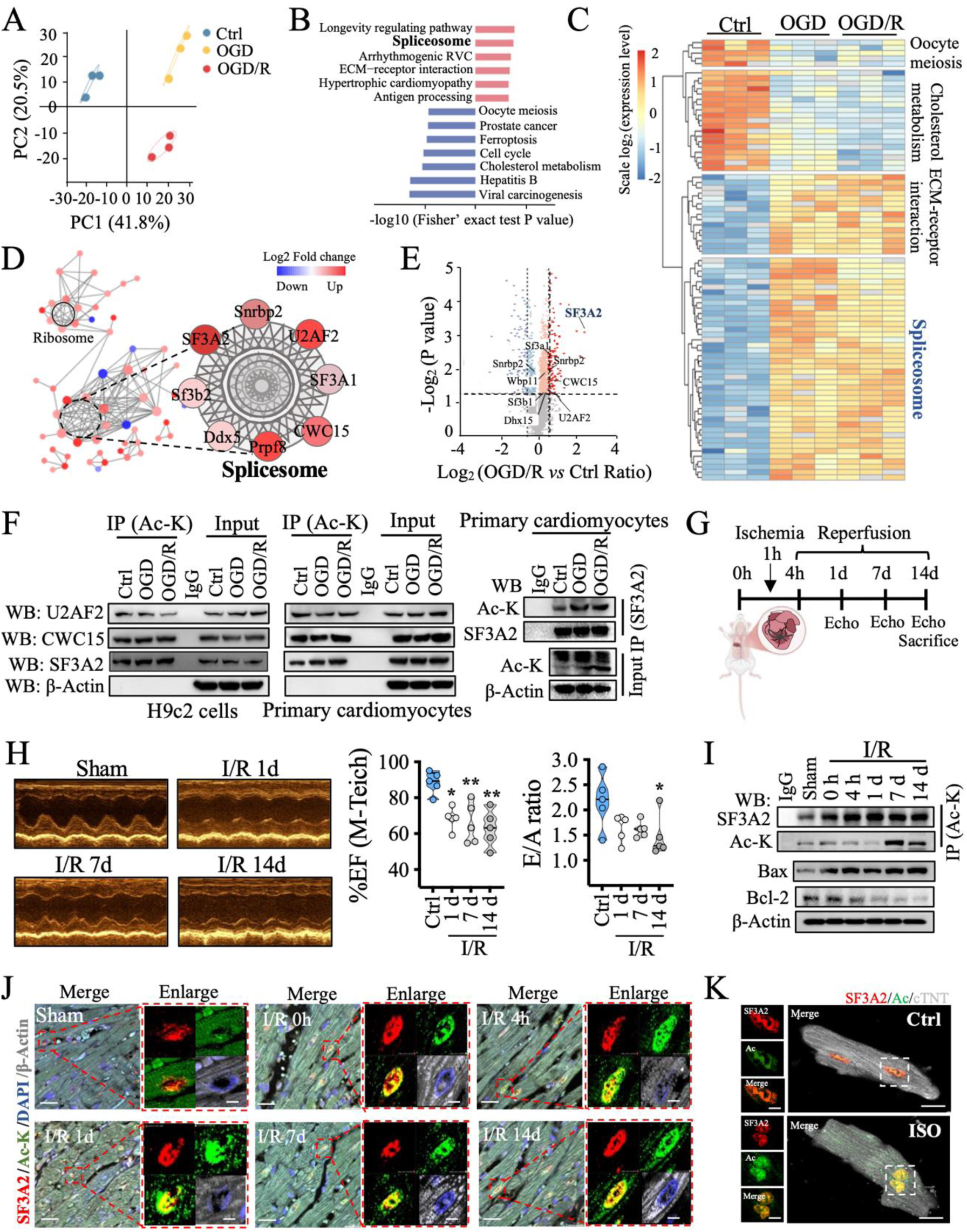
SF3A2 hyperacetylation in cardiomyocytes is a key regulator underlying pathological myocardial remodeling. **(A)** Principal component analysis (PCA) of acetylated proteins across control, oxygen/glucose deprivation (OGD), and OGD/reperfusion (OGD/R) groups. **(B)** KEGG pathway enrichment analysis of differentially acetylated proteins in OGD and OGD/R groups compared to the control. **(C)** Heatmap displays the significantly hyperacetylated proteins in the control, OGD, and OGD/R groups. **(D)** Protein-protein interaction (PPI) network analysis identifies central mediators of acetylation proteins in cardiomyocytes following OGD and OGD/R incubation. **(E)** Volcano plot analysis identifies the most significantly altered acetylated proteins in the OGD/R *vs* control group. **(F)** The acetylation of spliceosome-associated proteins was validated by co-immunoprecipitation (co-IP). N = 3 per group for **(A)**-**(F)**. **(G)** Schematic diagram of animal experiment design. A rat model of myocardial pathological remodeling was established by 1-hour myocardial ischemia followed by reperfusion for 4 hours, 1 day, 7 days, or 14 days. **(H)** Representative echocardiography and quantification of EF and E/A ratio at day 1, 7 and 14. **(I)** The immunoblots of acetylated SF3A2, Bax and Bcl-2. **(J)** Representative images of immunofluorescence (IF) staining of SF3A2, Ac-K, DAPI, and β-Actin in heart slices. Scale bar = 20 μm or 5 μm. **(K)** IF staining of Ac-SF3A2 and cTNT in adult mouse cardiomyocytes (ACMs). Scale bar = 20 μm or 5 μm. N = 5 per group for **(H)**-**(K)**. \**P* < 0.05 and \*\**P*< 0.01 by one-way ANOVA and Tukey multiple comparisons test. All pairwise comparisons were made.

### 2 The SF3A2 deacetylation at lysine 10 rescues mitochondrial dysfunction and myocardial hypertrophy induced by OGD/R and ISO incubation

Given the significance of SF3A2 acetylation in myocardial pathological remodeling, we next identify the specific acetylation sites to explore therapeutic strategies. Amino acid sequence analysis from acetylomic revealed that lysine (K) acetylation was strongly associated with a flanking glycine (G) at the preceding position and threonine (T) at the following position under OGD or OGD/R conditions (Figure 2A and Figure S3A). Conservation analysis indicated that the K10 residue within a Gly-Lys-Thr (G-K-T) motif of SF3A2 is highly conserved from Homo sapiens to Sus scrofa (Figure 2B). Additionally, LC-MS/MS analysis identified lysine 10 (K10) as the major acetylation site on SF3A2 (Figure 2C and Figure S3B). To further investigate the functional role of SF3A2, we generated a SF3A2 (K10) antibody that specifically recognizes acetylated SF3A2 at lysin 10 (Figure S3C-S3E). As shown in Fig. 2D, both OGD and OGD/R treatments induced the expression of acetylated SF3A2 (K10) in H9c2 and primary cardiomyocytes, with the OGD/R group exhibiting a higher level of acetylation than the OGD group. Furthermore, immunoblotting with a SF3A2 (K10) antibody confirmed that SF3A2 acetylation at lysin 10 increased in heart tissues following I/R injury in a time-dependent manner (Fig. 2E). Therefore, we generated stable cardiomyocyte lines using lentiviral infection and site-directed mutagenesis, expressing either an SF3A2 acetylation-deficient (K10R) or an SF3A2 acetylation-mimetic (K10Q) mutant, to investigate the role of SF3A2 (K10) acetylation in mitochondrial function and pathological remodeling (Fig. S4A). Mitochondrial membrane potential (MMP) and apoptosis results revealed that OGD/R incubation markedly increased depolarization of MMP and apoptosis in the negative control (NC), wild type (WT), and K10Q groups, whereas the K10R mutant effectively prevented these detrimental effects (Fig. 2F-2G). Additionally, the K10R mutant cardiomyocytes has higher basal oxygen consumption respiration (OCR), the capacity of maximal respiration and ATP production OCR than the WT group in response to OGD/R stress (Fig. 2H). To establish a direct link between SF3A2 acetylation and mitochondrial OXPHOS dysfunction in myocardial remodeling, we simultaneously assessed mitochondrial biogenesis, fission/fusion, and SF3A2 acetylation in mutant cardiomyocytes. Compared with the WT + ISO group, the K10R mutation rescued the ISO-induced mitochondrial fissions, SF3A2 hyperacetylation, and the reduction in mitochondrial fluorescence intensity (Fig. 2I and Fig. S4B). Moreover, correlation analysis demonstrated significant negative correlations between SF3A2 acetylation levels and both the mitochondrial fusion and mitochondrial biogenesis (Fig. 2J and Fig. S4C). Consistent with this, SF3A2(K10) deacetylation in ACMs also protected against MMP depletion in ISO-induced myocardial hypertrophy (Fig. 2K). Collectively, these results indicate that deacetylation of SF3A2 at K10 suppresses myocardial injury and hypertrophy by attenuating mitochondrial dysfunction.

**Fig. 2.**
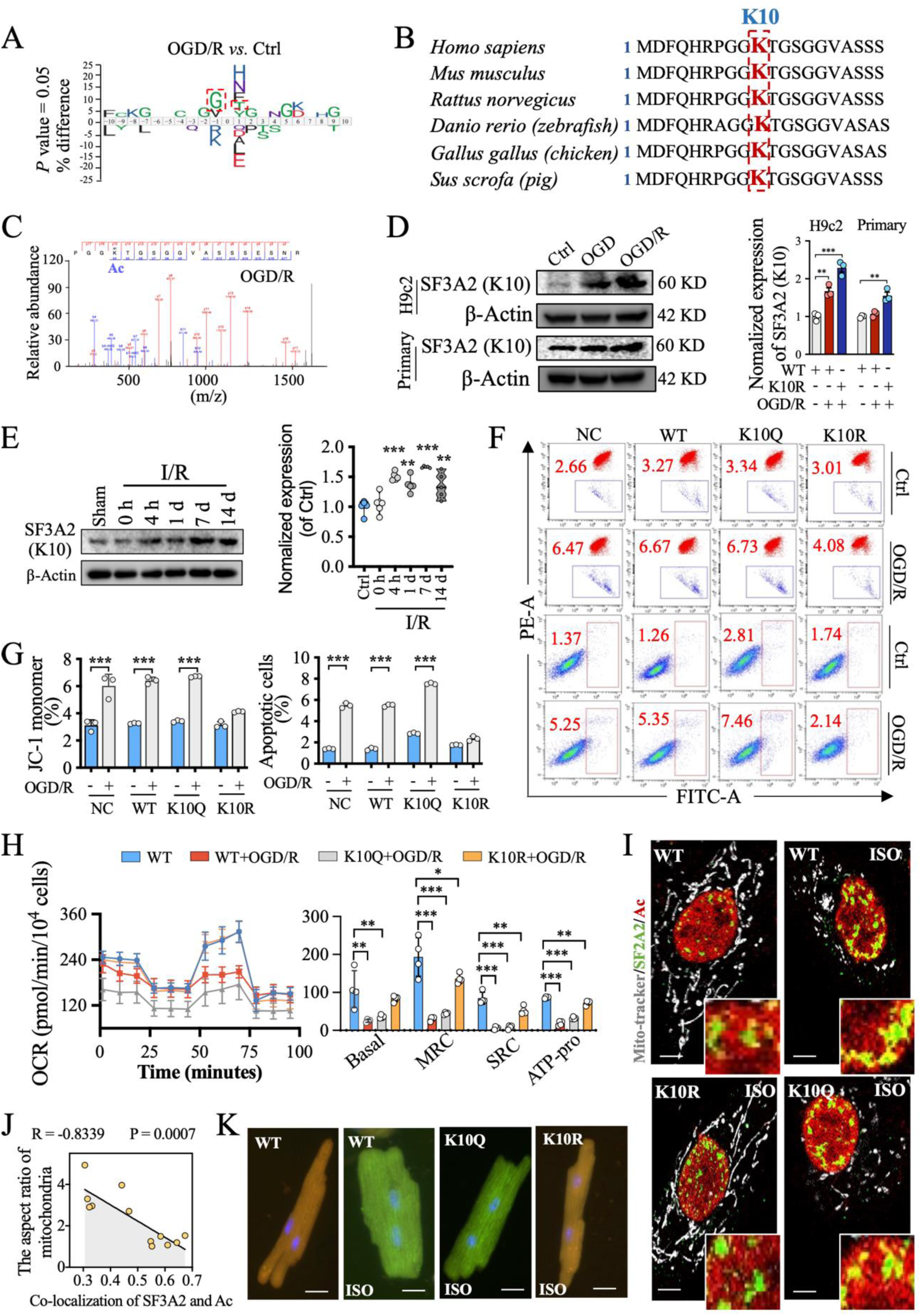
Deacetylation of lysine 10 on SF3A2 represents a key target for delaying myocardial pathological remodeling. **(A)** Amino acid sequence analysis from acetylomics in OGD/R *vs* control group. N = 3 per group. **(B)** Conservation analysis of the amino acid sequence on SF3A2 from Homo sapiens to Sus scrofa. **(C)** LC-MS/MS analysis was applied to identify the acetylated site on SF3A2 under OGD/R incubation. N = 3 per group. **(D)** Quantification of acetylation levels of SF3A2(K10) in H9c2 and primary cardiomyocytes. N = 3 per group. **(E)** Western blot and quantification of SF3A2(K10) acetylation in I/R-induced heart tissues. N = 5 per group. **(F-G)** Flow cytometry (FCM) was used to analyze the mitochondrial membrane potential (MMP) and apoptosis in SF3A2(K10) mutant cardiomyocytes. **(H)** The Seahorse XFe24 high-resolution respirometry was used to analyze basal oxygen consumption (Basal), capacity of maximal respiration (MRC), spare capacity of respiration (SRC) and ATP production oxygen consumption (ATP-pro) in the WT, K10R and K10Q cardiomyocytes. **(I)** Representative IF images of mitochondria, SF3A2 and Ac in mutant H9c2 cells. Scale bar = 5 μm. **(J)** Pearson correlation analysis of SF3A2 acetylation and mitochondrial fission. N = 3 per group for **(F)**-**(J)**. **(K)** Fluorescence MMP staining of ACMs. N = 5 per group. Scale bar = 20 μm. \**P* < 0.05, \*\**P*< 0.01 and \*\*\**P*< 0.001. Significant differences between groups were determined by one-way ANOVA followed by Tukey’s multiple comparison test.

### 3 Improvement of myocardial pathological remodeling after SF3A2(K10R) delivery

Given that SF3A2(K10R) prevented cardiomyocyte remodeling upon treatment with OGD/R and ISO in vitro, we performed the rescue experiment by injecting adenovirus carrying WT and SF3A2(K10R) into area of cardiac injury immediately after I/R in vivo. As shown in experimental workflow (Fig. 3A), AAV9 was chosen as the delivery system because it effectively infects the hearts of mice and rats. To ensure cardiomyocyte specificity, we used the cardiac troponin T (cTnT) promoter to drive SF3A2 expression. After I/R, AAV9-cTnT carrying WT and K10R was injected directly into the ischemic area (Fig. 3A-3B). As expected, immunohistochemical analysis confirmed that SF3A2(K10) acetylation was significantly elevated in the WT+I/R group compared to the WT group, and this increase was effectively suppressed in the K10R+I/R group (Fig. 3C). While heart function remained a comparable decline in systolic function in both WT+I/R and K10R+I/R groups at 2 weeks post-I/R, the K10R+I/R group exhibited a progressive improvement in systolic function from week 4 to week 8, compared to the WT+I/R group (Fig. 3D and Fig. S5). Accordingly, histopathological analysis at 8 weeks post-AAV injection revealed that SF3A2(K10) deacetylation alleviated I/R-induced aggravation of disturbed myofibrillar arrangement, myofibrillar swelling and myocardial fibrosis, as evidenced by H&E and Masson staining (Fig. 3E). There was no significant difference in histopathological phenotype between the NC and WT groups (Fig. 3E). Furthermore, WGA staining indicated a marked reduction in cardiomyocyte hypertrophy in the K10R+I/R group compared to the WT+I/R group (Fig. 3E). Moreover, transmission electron microscopy (TEM) analysis revealed that the mutation at lysine 10 significantly attenuated I/R-induced mitochondrial damage, including reduced mitochondrial vacuolization, loss of cristae, and decreased mitochondrial number (Fig. 3F). As shown in Figure 3G, the WT+I/R hearts exhibited a significantly decreased Bcl-2/Bax ratio, which was accompanied by increased acetylation of SF3A2 at lysine 10. In contrast, deacetylation of this site conversely led to an elevated Bcl-2/Bax ratio. These results provide compelling evidence that deacetylation of SF3A2 at lysine 10 alleviates mitochondrial dysfunction, restores heart function, and attenuates pathological remodeling in *in vivo* cardiomyocytes.

**Fig. 3.**
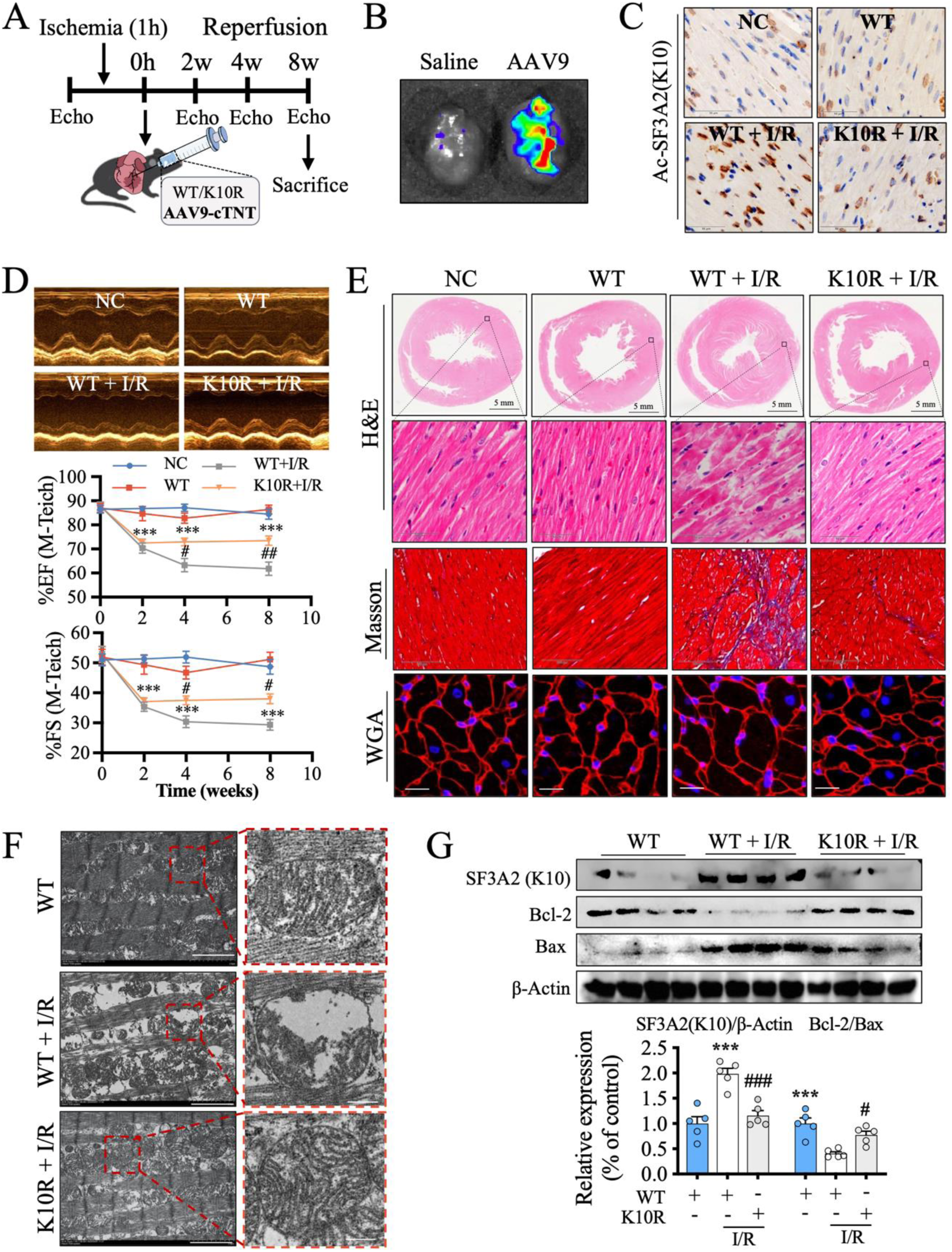
AAV9-cTNT mediated SF3A2(K10R) delivery attenuates myocardial hypertrophy, fibrosis, and dysfunction. **(A)** Schematic diagram of experiment protocol. Rats were randomly infected AAV9-SF3A2 (WT) or AAV9-SF3A2(K10R) via direct injection into the ischemic injury area, followed by subsequent analyses. **(B)** Fluorescence imaging of heart confirmed the precise delivery and transduction efficiency of AAV9 following targeted intramyocardial injection. **(C)** Immunohistochemistry (IHC) staining of Ac-SF3A2(K10). Scale bar = 50 μm. **(D)** Echocardiographic analyses of EF and FS. **(E)** Representative images of H&E (scale bar = 5 mm or 50 μm), Masson (scale bar = 100 μm), and WGA (scale bar = 20 μm) staining. **(F)** Representative images of transmission electron microscope (TEM) in heart tissues. Scale bar = 2 μm or 500 nm. **(G)** Western blot analysis of the acetylated SF3A2 (K10), Bax and Bcl-2 expression. N = 5 per group for **(A)**-**(G).** \*\*\**P* < 0.001, compared to WT group, ^#^*P* < 0.05 and ^###^*P* < 0.001, compared to WT+I/R group. Analyses performed by one-way ANOVA and Tukey multiple comparisons test. All pairwise comparisons were made.

### 4 SF3A2 hyperacetylation drives cardiomyocyte lipotoxicity, a key mechanism implicated in myocardial pathological remodeling

Given that pathological remodeling involves the alterations in mitochondrial energy metabolism, we sought to define the mechanistic role of SF3A2 through untargeted metabolomics and targeted lipidomic. Following the experimental workflow, plasma and left ventricular tissues were collected from WT and K10R hearts following I/R injury to identify the characteristically metabolic accumulation (Fig. 4A). Untargeted metabolomics revealed distinct clustering of plasma and heart tissue metabolites from WT, WT+I/R and K10R+I/R groups by PCA (Fig. 4B). Volcano plot analysis of differential metabolites revealed that I/R injury resulted in 57 upregulated and 1399 downregulated metabolites in plasma (Fig. S6A). In contrast, SF3A2 deacetylation in I/R-induced myocardial pathological remodeling demonstrated a substantial reversal, with 1496 upregulated and 60 downregulated metabolites in plasma (Fig. 4C). In heart tissues, the WT+IR group exhibited 245 upregulated and 169 downregulated metabolites (Fig. S6A). In comparison, the K10R+IR group showed 155 upregulated and 191 downregulated metabolites (Fig.4C). To investigate the correlation between plasma and heart tissue metabolites, we identified overlapping differential metabolites using a Venn diagram and subsequently performed enrichment analysis (Fig.4D). The 121 enriched metabolites were subjected to KEGG pathway analysis, which indicated that SF3A2 deacetylation primarily regulates metabolites associated with lipid metabolism and central carbon metabolism in both heart tissues and plasma (Fig.4D and Fig. S6B-S6D). In accordance with the changes in metabolomics, targeted lipidomic profiling revealed that SF3A2(K10R) reverses the I/R-induced accumulation of free fatty acids (FFAs) and sphingomyelin (SM), along with the reduction in triacylglycerols (TAG) in plasma (Fig. S7A-S7B). In heart tissues, I/R injury primarily led to the accumulation of ceramide (Cer), cholesteryl ester (CE), and phosphatidylinositol (PI), which was significantly attenuated by SF3A2(K10R) treatment (Fig. S7B-S7C). Impaired OXPHOS compromises fatty acid β-oxidation, leading to the accumulation of FFAs and their cytotoxic intermediates (e.g., acetyl-coenzyme A, CE, and Cer), thereby inducing lipotoxicity that impairs heart function^12^. Our correlation analyses of lipidomic and echocardiographic data revealed that plasma levels of FFAs and SM, as well as myocardial levels of Cer, CE and PI, were inversely correlated with the heart ejection fraction (Fig. 4E and Fig. S8). We therefore hypothesized that deacetylation of SF3A2 attenuates fatty acid uptake, thereby ameliorating lipotoxicity. To test this, we measured the expression and translocation of CD36 (a primary fatty acid transporter), the expression of ATP-citrate lyase (ACLY, a rate-limiting enzyme for de novo lipogenesis), and the levels of acetyl-CoA (a central metabolite in fatty acid catabolism). Western blotting and immunofluorescence analysis revealed that SF3A2 deacetylation do not alter the total protein expression of CD36 but rather reduce its translocation from the cytoplasm to the plasma membrane in cardiomyocytes (Fig. 4F-4G). The protein expression of ACLY was inhibited by the deacetylation of SF3A2 (Fig. 4F). Moreover, the deacetylation of lysine 10 in SF3A2 also alleviated the accumulation of acetyl-CoA induced by pathological remodeling (Fig. 4H). Collectively, these results illustrated that SF3A2 deacetylation reverses a pathological cycle during myocardial remodeling: upon reperfusion following ischemia, cardiomyocytes exhibit a substantial uptake of FFAs from the blood. However, impaired OXPHOS disrupts FFAs β-oxidation, leading to the accumulation of FFAs and cytotoxic intermediates (acetyl-CoA, CE, and Cer). This lipid overload induces lipotoxicity, triggering cardiomyocyte apoptosis and ultimately impairing heart function. Concurrently, the apoptotic process activates sphingomyelinase, which hydrolyzes membrane sphingomyelin (SM) to generate ceramide (Cer) (Fig. 4I). Therefore, these results indicate that the protective effect of SF3A2 deacetylation against myocardial pathological remodeling is mediated through the suppression of CD36-mediated lipotoxicity.

**Fig. 4.**
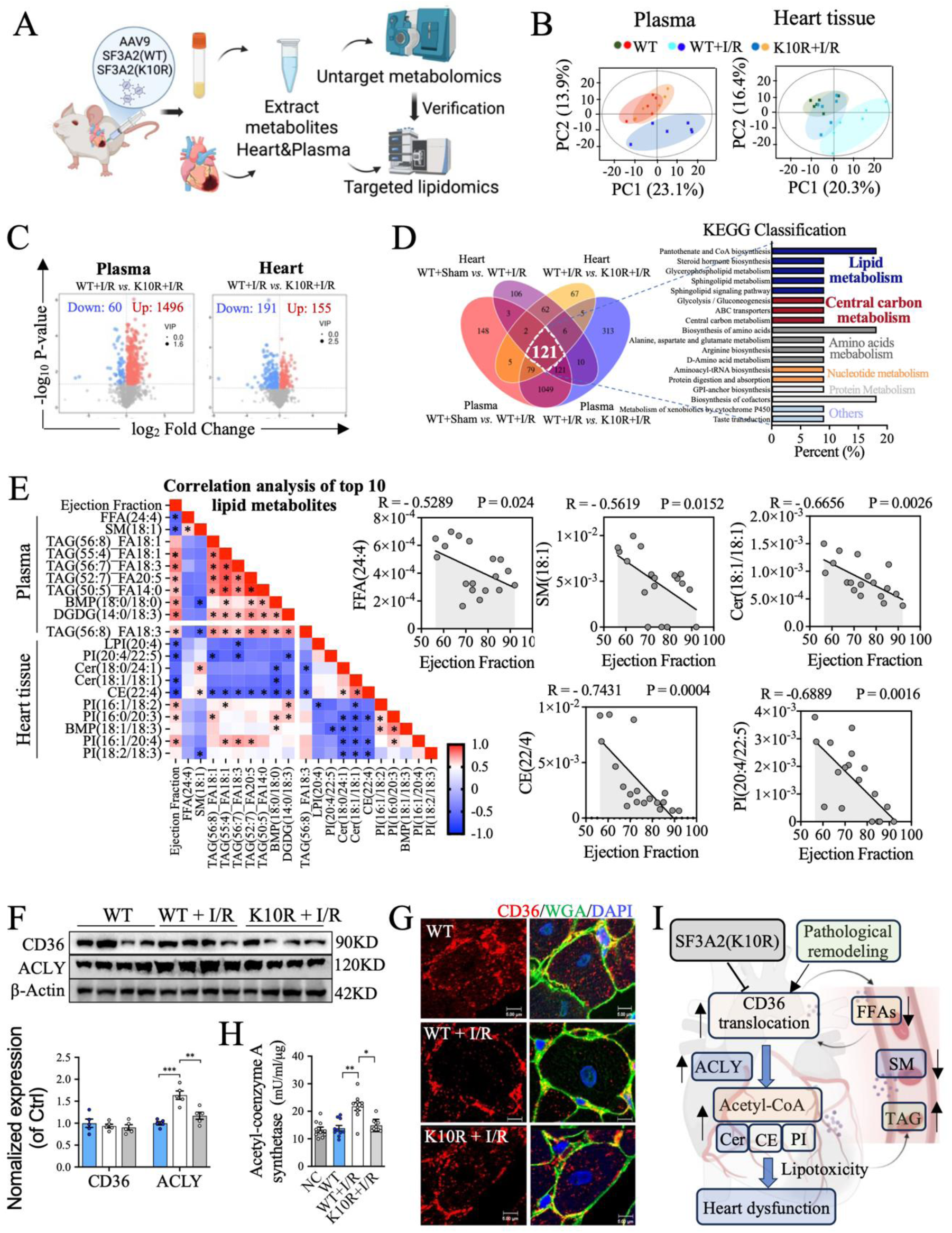
SF3A2 hyperacetylation promotes cardiomyocyte lipotoxicity. **(A)** Schematic diagram of targeted and untargeted metabolomics analysis in cardiac tissue and plasma. **(B)** PCA reveals distinct clustering of cardiac and plasma metabolites among groups WT, WT +I/R, and K10R+I/R. **(C)** Volcano plot analysis of differential lipid metabolites in cardiac tissue and plasma. **(D)** The overlapping differential metabolites was enriched using a Venn diagram analysis in WT *vs.* WT+I/R and WT+I/R *vs.* K10R+I/R groups. **(E)** Correlation analysis of lipid metabolites and heart function. **(F)** Western blot analysis of CD36 and ACLY. **(G)** Immunofluorescence staining of CD36 in cardiac tissue sections. Scale bar = 5 μm. N = 5 per group for **(B)-(G)**. **(H)** Assessment of acetyl-CoA activity in cardiac tissue. N = 9 per group. **(I)** Schematic representation of CD36-mediated lipotoxicity in pathological cardiac remodeling. Significant differences between groups were determined by one-way ANOVA followed by Tukey’s multiple comparison test. \**P* < 0.05, \*\**P* < 0.01, \*\*\**P* < 0.001.

### 5 Deacetylation of SF3A2 at lysine 10 promotes the aerobic oxidation of glucose in cardiomyocytes

Cardiomyocytes primarily utilize glucose and fatty acids as metabolic substrates for energy production through TCA cycle and OXPHOS. Given that our lipidomic data demonstrating that SF3A2 deacetylation suppresses fatty acid uptake, we hypothesize a compensatory shift toward enhanced glucose metabolism in cardiomyocytes. Therefore, a ^13^C carbon tracing method based on LC-MS was employed to map the fate of glucose metabolism mediated by SF3A2 (Fig. 5A). Based on the labeling patterns of glucose metabolites, PCA demonstrated distinct metabolic fluxes in glycolysis and the TCA cycle among the WT, WT+OGD/R, and K10R groups (Fig. 5B). In contrast, fluxes in the pentose phosphate pathway (PPP) were largely comparable among these groups (Fig. 5B). As shown in Fig. 5C-5D, the levels of several glycolysis metabolites (F1,6P, GAP, 3PG, G3P, Ser, Pyr and Lac) from K10R+OGD group were decreased, compared to WT+OGD group, suggesting an inhibitory effect on glycolysis. Mutant SF3A2(K10R), but not the WT H9c2 cells, efficiently reduced the level of 6PG,E4P, R5P and another major PPP product, S7P (Figure 5E). We also performed Seahorse assays and found that the OGD/R-triggered glycolysis was markedly attenuated by SF3A2 deacetylation (Figure 5F-5G). In addition, glucose flux into the TCA cycle was completely increased in K10R+OGD/R group (Fig. 5H-5I). Citrate serves as the first intermediate and a key regulatory node in the TCA cycle, linking carbohydrate metabolism to lipid biosynthesis^23^. Under OGD/R conditions, a concurrent increase in unlabeled citrate and decrease in the mass (M)+2 citrate were observed, suggesting an accumulation of non-oxidized citrate (Fig. 5J). In contrast, SF3A2(K10R) led to a marked increase in mass (M)+2 citrate, indicating an accelerated TCA cycle flux (Fig. 5J). Additionally, SF3A2 deacetylation elevated ATP synthesis in pathological remolding cardiomyocytes (H9c2 cells, ACMs, and rat models) (Fig. 5K). Collectively, these results indicated that SF3A2 deacetylation redirects glucose metabolism from anaerobic glycolysis into the TCA cycle.

**Fig. 5.**
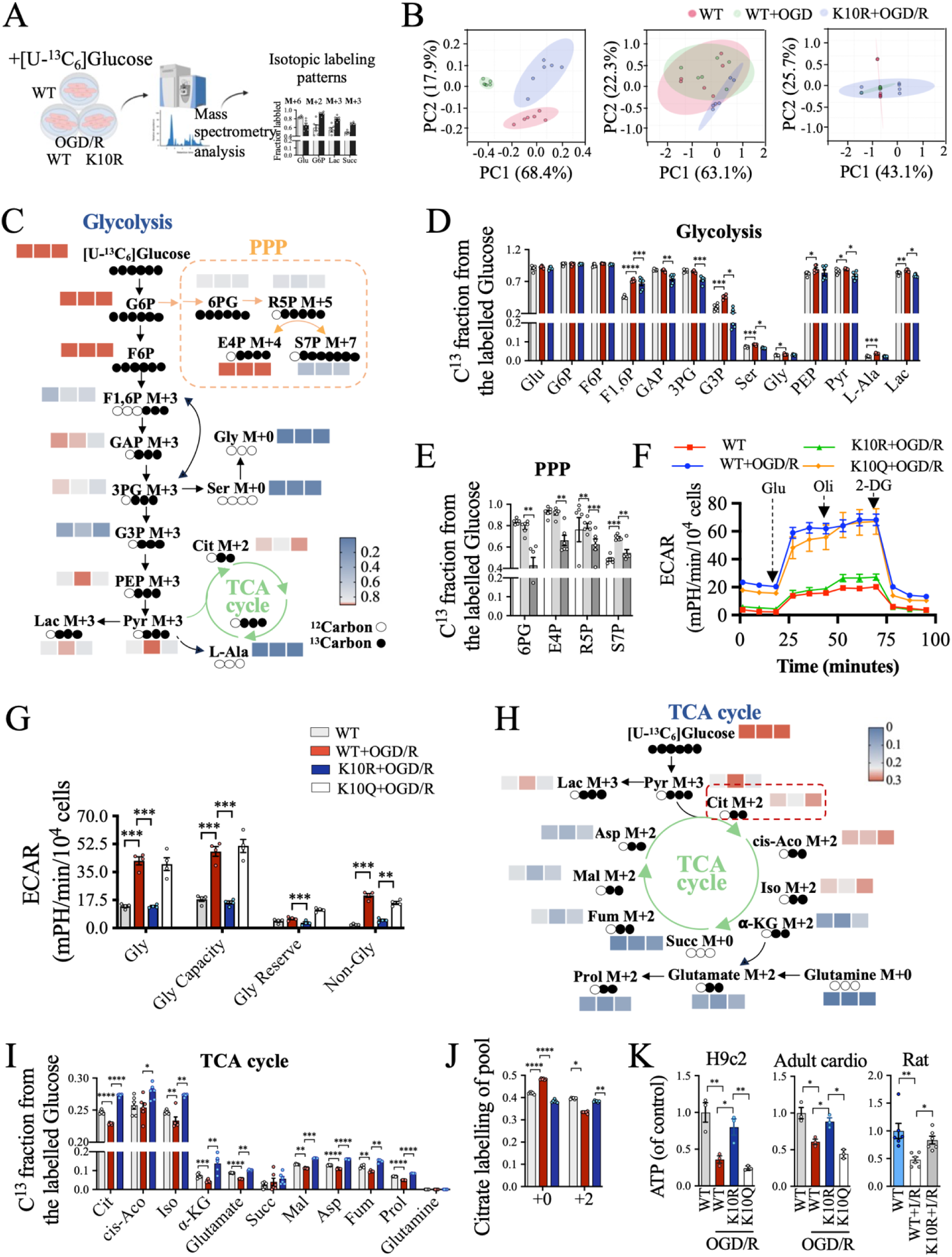
SF3A2 deacetylation shifts myocardial metabolism from anaerobic glycolysis toward the TCA cycle in response to I/R stress. **(A)** Schematic diagram illustrating the U-¹³C₆-glucose tracing analysis using LC-MS in SF3A2 mutant cardiomyocytes. **(B)** PCA demonstrates the distinct clustering of ^13^C fraction from the labelled glucose. **(C)** The conversion pathway of U-^13^C6-glucose into glycolysis– and pentose phosphate pathway (PPP)-related metabolites. **(D-E)** The quantification of ^13^C fraction from the labelled glucose. **(F-G)** The Seahorse XFe24 high-resolution respirometry was used to analyze the basal glycolysis (Gly), glycolysis capacity (Gly Capacity), glycolysis reserve (Gly Reserve) and non-glycolysis acidification (Non-Gly) in mutant cardiomyocytes. **(H)** The TCA cycle pathway of U-^13^C6-glucose labeled metabolites. **(I)** The quantification of ^13^C fraction from the labelled glucose of TCA cycle. **(J)** Quantification of (M)+0 citrate and (M)+2 citrate. **(K)** The content of ATP in cardiomyocytes and heart tissues. N = 3 per group for **(B)**-**(K)**. \**P* < 0.05, \*\**P* < 0.01 and \*\*\**P* < 0.001 by one-way ANOVA and Tukey multiple comparisons test. All pairwise comparisons were made.

### 6 SF3A2 deacetylation attenuates myocardial pathological remodeling through regulating the alternative splicing of Nnt to preserve mitochondrial function

SF3A2 is an essential splicing factor in the U2 snRNP complex that exerts its function by regulating the AS of key downstream genes^18^. To systematically identify its downstream targets, we conducted full-length transcriptome sequencing and AS analysis in hearts. The volcano plot analysis identified a total of 2,656 differentially expressed genes (DEGs) in the K10R+I/R group compared to the WT+I/R group, of which 1,519 were upregulated and 1,137 were downregulated (Fig. 6A). KEGG and heatmap analysis on these DEGs revealed that SF3A2 deacetylation upregulated the gene classes involved in OXPHOS, cardiac muscle contraction, carbon metabolism and TCA cycle and downregulated of gene classes involved in focal adhesion, protein digestion and absorption and ECM-receptor interaction, consistent with the lipidomic and metabolic flux observations (Fig. 6B-6C). AS profiles by RNA-seq analysis identified that AS events regulated by SF3A2 deacetylation included skipped exons (SE), the most abundant category (36%), followed by alternative 5′ splice sites (A5SS) (18%) and alternative 3′ splice sites (A3SS) (14%) (Fig. 6D). To determine the subset of DEGs that were driven by AS, we performed an overlap analysis between the DEGs and differentially AS. As shown in Figure 6E, we enriched a list of 30 overlapping genes and validated the mitochondria function-related gene, including Uqcrb, Tango2, Nnt and Lyplal1. qPCR analysis revealed that pathological remodeling in cardiomyocytes downregulated the expression of Uqcrb, Tango2 and Nnt but upregulated Lyplal1 (Fig. 6F). In contrast, SF3A2 deacetylation specifically enhanced the expression of Nnt (Fig. 6F). Percent Spliced In (PSI) analysis revealed that specifically skipping of exon 2 in the Nnt transcript (Nnt-Δe2) in response to I/R stress (Fig. 6G). Conversely, deacetylation of lysine 10 on SF3A2 promoted the retention of this specific exon, suggesting SF3A2 deacetylation modulates the AS of Nnt (Fig. 6G). Nnt catalyzes NAD^+^ regeneration in cardiomyocytes, sustaining mitochondrial respiration and redox homeostasis essential for heart function^24^. We therefore assessed NAD⁺ levels in *in vivo* and found that SF3A2 deacetylation promotes myocardial NAD⁺ level in response to I/R stress (Fig. 6H). To investigate the functional significance of Nnt-Δe2, we employed siRNA-mediated Nnt knockdown in site-specific mutant H9c2 cells, followed by an assessment of OXPHOS function. As expected, SF3A2 deacetylation restored basal OCR, the capacity of maximal respiration and ATP production OCR in si-Ctrl cells; however, this effect was abolished by Nnt knockdown (Fig. 6I). Our study reveals that SF3A2 deacetylation modulates the AS of Nnt-Δe2, thereby attenuating mitochondrial dysfunction to mitigate pathological remodeling.

**Fig. 6.**
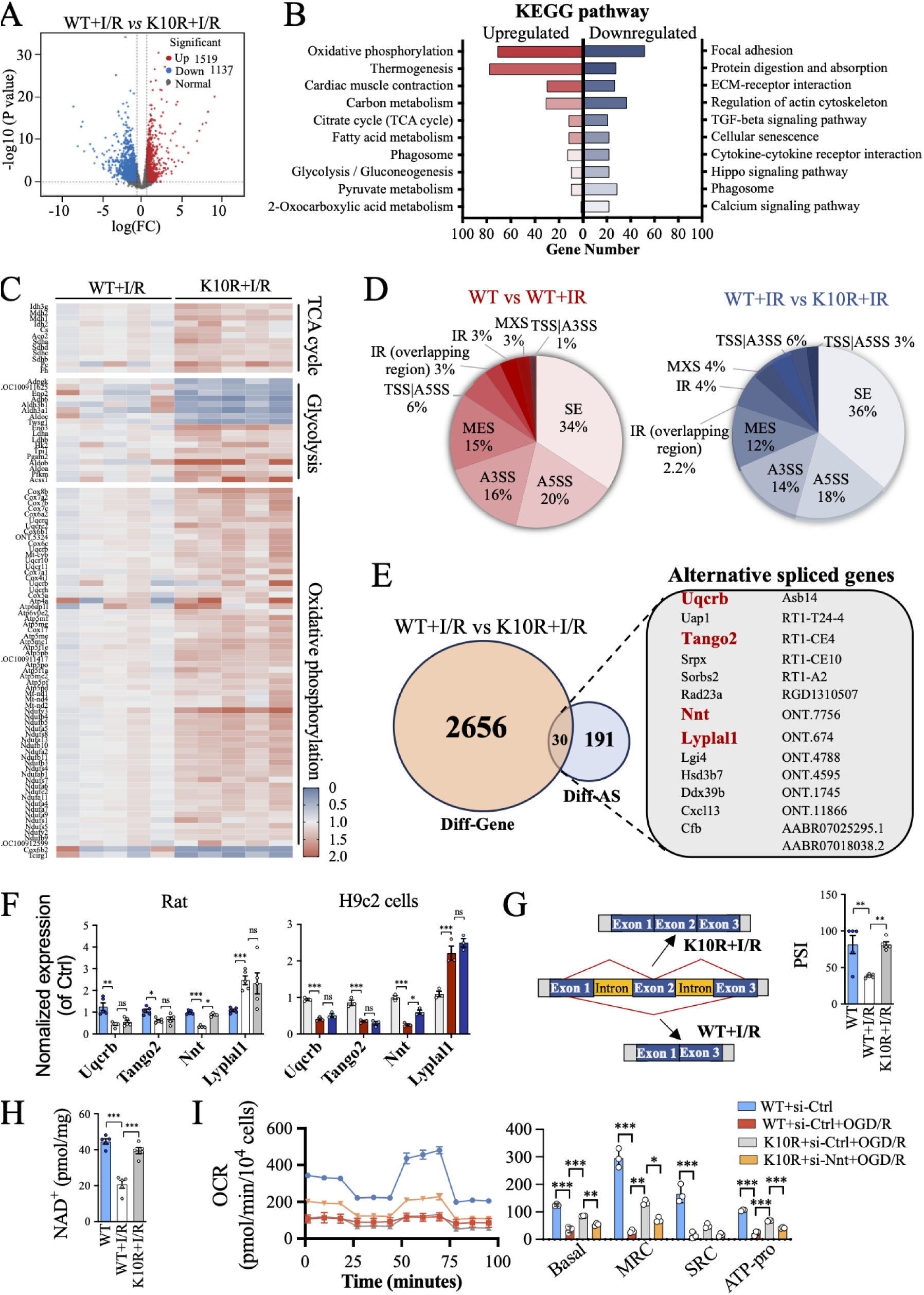
SF3A2 hyperacetylation induces exon 2 skipping in the Nnt transcript through. **AS. (A)** Volcanic plot illustrating differentially expression genes (DEGs). Red represents up-regulated, and blue represents down-regulated. **(B)** KEGG enrichment analysis of DEGs. **(C)** The heatmap shows the DEGs regulated by SF3A2 deacetylation. **(D)** The percentage of different splicing events in the AS of differentially expressed gene. Exon skipping (SE), alternative 5′ splice site (A5SS), alternative 3′ splice site (A3SS), mutually exclusive exons (MES) and intron retention (RI). **(E)** Genes enriched by Venn diagrams that undergo both AS and differential transcription. N = 5 per group for **(A)**-**(E)**. **(F)** qPCR analysis was use to validated the differentially expressed genes in heart tissues and cardiomyocytes. N = 5 per group for heart tissues. N = 3 per group for cardiomyocytes. **(G)** Schematic diagrams (left panel) showing detailed splicing sites of Nnt genes. The right panel quantifies exon 2 skipping in Nnt using percent-spliced-in (PSI) analysis. **(H)** Quantification of NAD^+^ levels in heart tissues. N = 5 per group for **(G)**-**(H)**. **(I)** Assessment of mitochondrial OXPHOS in Nnt knockdown cardiomyocytes using a Seahorse XFe24 analyzer. N = 3 per group. \**P* < 0.05, \*\**P* < 0.01 and \*\*\**P* < 0.001. All pairwise comparisons were made. Statistical comparisons between groups were analyzed by one-way ANOVA, followed by Tukey’s post hoc test.

### 7 SIRT7 binds to SF3A2 and influences its deacetylation function

To identify upstream pathway of SF3A2, IP-MS was performed to identify the interacting partners of SF3A2 (Fig. 7A). Among the ten identified proteins, three proteins associated with heart function were selected for further validation (Fig. 7B). Co-IP assays verified that SF3A2 and Ddx1 form a complex under OGD/R stress (Fig. 7C). Moreover, the co-localization of SF3A2 and Ddx1 in cardiomyocytes was abolished by SF3A2 deacetylation in pathological remodeling (Fig. 7D). Ddx1, an RNA helicase in cardiomyocytes, resolves secondary structures to ensure efficient mRNA processing and translation, thereby supporting mitochondrial biogenesis and cell survival under metabolism stress^25^. Given that the SF3A2-Ddx1 complex is implicated in mRNA processing, we asked if SF3A2 acetylation is regulated by acetyltransferases or deacetylases. To investigate the regulators of SF3A2 acetylation, we employed shRNA-mediated knockdown of acetyltransferases and deacetylases in SF3A2-mutant H9c2 cells. The immunoblot experiments showed that the acetylation of SF3A2 was mainly regulated by SIRT7 (Fig. 7E). We next sought to determine the function of SIRT7 in acetylation/deacetylation of SF3A2. In NC cells, SF3A2 deacetylation suppressed OGD/R-induced mitochondrial fission and impaired biogenesis, whereas this effect was markedly attenuated in SIRT7-deficient cardiomyocytes (Fig. 7F-7G). Moreover, SIRT7 knockdown abolished the ability of SF3A2 deacetylation to attenuate OGD/R-induced apoptosis (Fig. 7H and Fig. S9A). Taken together, these results demonstrate that impaired SIRT7 function results in SF3A2 hyperacetylation in myocardial pathological remodeling.

**Figure 7.**
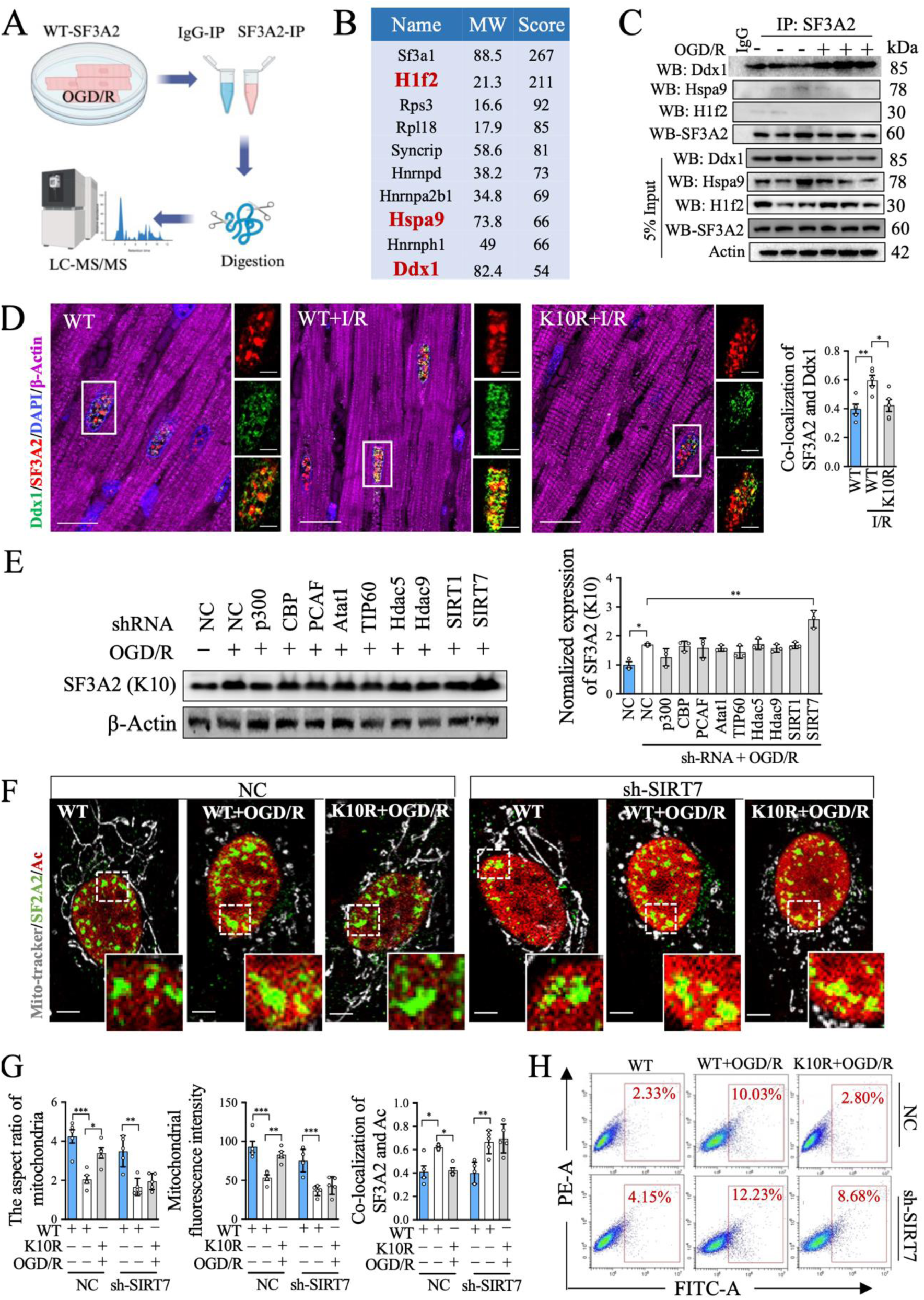
SIRT7 binds to SF3A2 to influence its deacetylation function. (**A**) Immunoprecipitation coupled with mass spectrometry (IP-MS) was employed to identify the SF3A2-binding proteins. N = 3 per group. **(B)** Top 10 SF3A2-binding proteins ranked by binding score. **(C)** Co-IP assays were performed to identify SF3A2-interacting proteins. N = 3 per group. **(D)** Representative IF images and quantitative analysis of SF3A2 and Ddx1 co-localization in heart slices. Scale bar = 5 μm or 20 μm. N = 5 per group. **(E)** Western blot assay and corresponding quantification of SF3A2(K10) acetylation in cardiomyocytes. **(F)** Representative IF images of mitochondria, SF3A2 and Ac in SIRT7 knockdown cardiomyocytes. Scale bar = 5 μm. **(G)** Quantification of the aspect ratio of mitochondria, mitochondrial fluorescence intensity, and the co-localization of SF3A2 and Ac in mutant cardiomyocytes. **(H)** FCM was used to analyze the apoptosis level in NC or SIRT7 knockdown cardiomyocytes. N = 3 per group for **(E)-(H)**. \**P* < 0.05, \*\**P* < 0.01 and \*\*\**P* < 0.001 by one-way ANOVA and Tukey multiple comparisons test. All pairwise comparisons were made.

### 8 Pharmacological activator of SIRT7 via small-molecule compounds offers a therapeutic strategy

Our findings demonstrated that AAV9-delivered SF3A2 deacetylation mitigates myocardial pathological remodeling and improves heart function. However, its clinical translation remains a future challenge. Natural saponins, as promising SIRTs activators, exhibit potential in mitigating cardiovascular pathologies by enhancing mitochondrial function and suppressing oxidative stress^26–28^. Based on the SIRT7 binding affinity, SF3A2 acetylation, NAD⁺ content, and MMP, we screened a library of small-molecule saponins to identify candidate natural compounds capable of activating the SIRT7–SF3A2 pathway in ACMs and H9c2 cells (Fig. 8A). This screen results revealed that resveratrol, 2-APQC, Rb1, Rb2, Rc, CK, and Rg1 could counteract the reduction in NAD^+^ and the depolarization of MMP induced by either OGD/R or ISO (Fig. S9B-S9D). As shown in Fig.8B and Fig.S10A, surface plasmon resonance analysis revealed that 2-APQC and ginsenoside Rg1 bind to the SIRT7 protein with KD values of 2.66 µM and 5.20 µM, respectively. The Cellular Thermal Shift Assay (CETSA) results further confirmed that ginsenoside Rg1 displays a stronger binding affinity with SIRT7, compared to 2-APQC (Fig. 8C). These results suggest that Rg1 binds to SIRT7 to deacetylate SF3A2, and ultimately rescues mitochondrial dysfunction. Thus, we further validated the cardioprotective function of Rg1 in *in vivo*. we treated mice with Rg1 concurrently with ISO subcutaneous injection to investigate its potential to attenuate pathological remodeling. Metoprolol was used as a positive control for cardioprotection. Echocardiographic results demonstrated that an 8-weeks ginsenoside Rg1 treatment significantly increased both the left ventricular ejection fraction and decreased the E/e’ ratio compared to the ISO group, indicating enhanced heart systolic and diastolic function (Fig. 8D-8E). The ratio of heart weight/tibia length and WGA staining results consistently demonstrated that Rg1 alleviated ISO-induced myocardial hypertrophy (Fig. 8F-8G and Fig. S10B). Furthermore, Rg1 treatment significantly rescued the ISO-induced depletion of ATP content in heart tissues (Fig. 8H). The increases in acetyl-CoA and ACLY expression, observed in ISO-treated mice were also reversed by Rg1, indicating that Rg1 reduces fatty acid uptake, thereby alleviating ISO-induced lipotoxicity (Fig. 8H-8I). To determine whether the amelioration of fatty acid uptake and mitochondrial dysfunction by Rg1 is directly dependent on SIRT7, we assessed CD36 translocation and MMP in ACMs. Fluorescence analysis revealed that the protective effects of Rg1 against ISO-induced CD36 translocation and mitochondrial depolarization were abolished by SIRT7 knockdown (Fig. 8J). Taken together, our findings indicate that pharmacological activation of the SIRT7-SF3A2 pathway attenuates myocardial pathological remodeling and improves heart function by suppressing lipotoxicity and restoring mitochondrial function.

**Figure 8.**
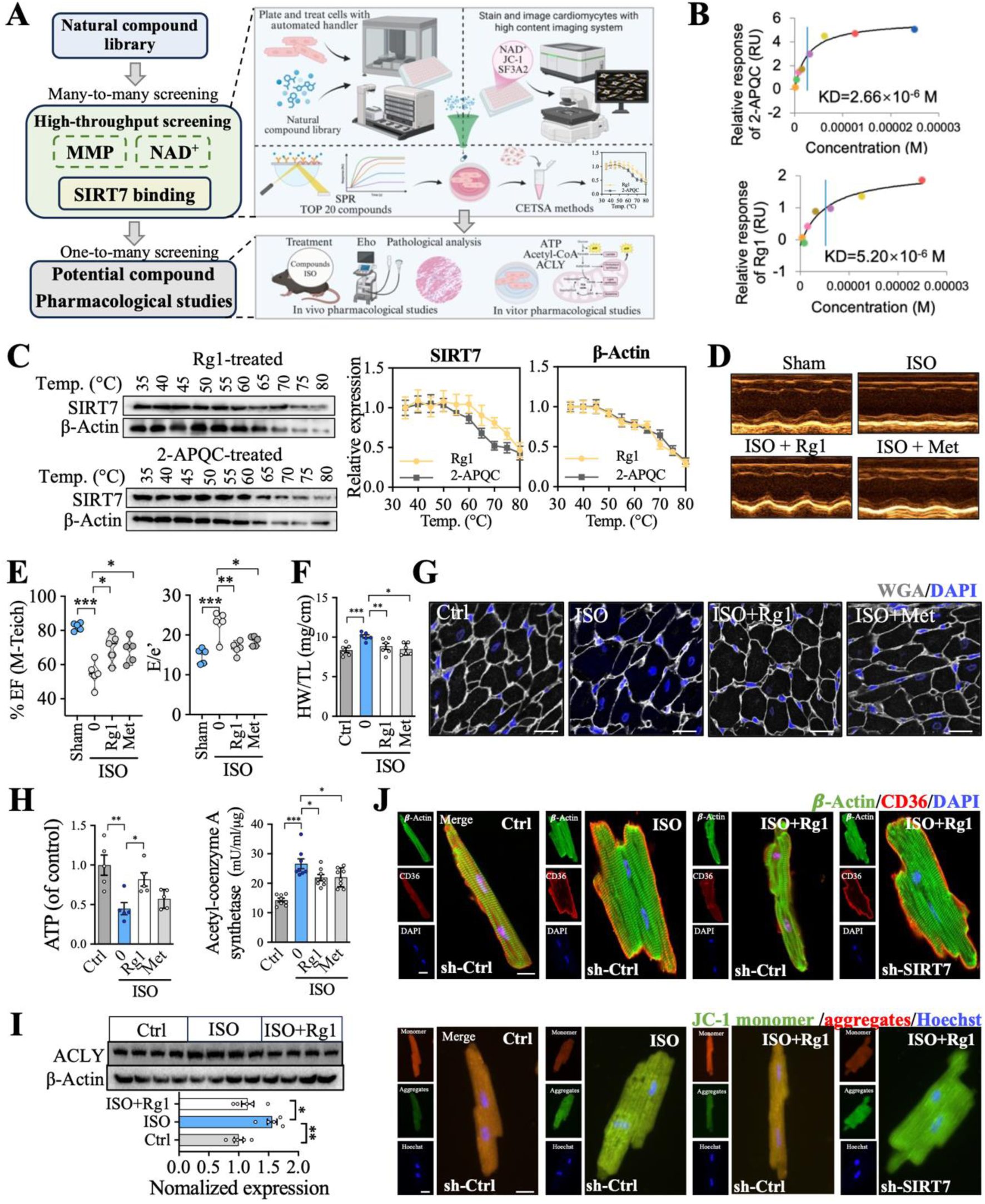
Identification of ginsenoside Rg1 as a lead compound targeting SIRT7 to suppress ISO-induced pathological remodeling. **(A)** Schematic diagram of high-throughput screening with NAD^+^-dependent SIRT7 activity in ACMs and H9c2. **(B)** Surface plasmon resonance (SPR) was used to analyze the binding between different small-molecule saponins and SIRT7. **(C)** The binding affinity of candidate small molecules to SIRT7 was assessed using the cellular thermal shift assay (CETSA). N = 3 per group for **(B)**-**(C)**. **(D)** Cardiac echocardiography was used to analyze the effect of ginsenoside Rg1 on ISO-induced pathological remodeling. Metoprolol (Met) as a positive control. **(E)** Quantification of left ventricular EF and E/e’ ratios. **(F)** The ratio of heart weight to tibia length (HW/TL) was quantified. **(G)** Representative images of WGA staining. Scale bar = 20 μm. **(H)** ATP and acetyl-coenzyme A levels were measured in the left ventricular tissue of mice. **(I)** Western blot and quantification of ACLY expression in heart tissues. **(J)** Representative IF images of CD36 in ACMs. Scale bar = 20 μm or 5 μm (zoom in). N = 5 per group for **(D)**-**(J)**. \**P* < 0.05, \*\**P* < 0.01 and \*\*\**P* < 0.001. Significant differences between groups were determined by one-way ANOVA followed by Tukey’s multiple comparison test.

## Discussion

Although intrinsic metabolic reprogramming of cardiomyocytes during pathological remodeling are well documented, these changes are often viewed as a compensatory or merely coincidental phenomenon rather than a cause of the disease, and thus have rarely been considered as a viable target for treatment^29, 30^. We set out to test the hypothesis that, instead, these metabolic reprogramming play a causal role in driving myocardial pathological remodeling. Consistent with this hypothesis, we identified a tight coupling between heart dysfunction and metabolic reprogramming in a rodent model of pathological remodeling. This reprogramming was driven by enhanced anaerobic glycolysis and intracellular lipid accumulation, and repressed fatty acid β-oxidation. The majority of mitochondrial ATP production in cardiomyocytes is derived from the oxidation of fatty acids and glucose^31^. Upon uptake into cardiomyocytes—a process partly mediated by CD36 and fatty acid transport protein-1—fatty acids are subjected to β-oxidation, thereby producing acyl-CoA for the TCA cycle^10^. The majority of pyruvate from glycolysis also is converted to acetyl-CoA by aerobic oxidation of glucose to anaplerosis that replenishes TCA intermediates^32^. In our study, we observed that in response to I/R or ISO challenge, CD36 translocate to the cardiomyocyte membrane, enhancing fatty acid uptake from the plasma. However, impaired mitochondrial OXPHOS function and citrate accumulation inhibit further β-oxidation of these fatty acids, leading to the accumulation of acyl-CoA. These findings indicate that myocardial remodeling drives a shift in metabolic substrate utilization toward glycolysis, compensating for reduced ATP generation from fatty acids. Importantly, targeted deacetylation of SF3A2 in left ventricular cardiomyocytes, achieved via AAV9-mediated gene delivery, restored heart systolic function by mitigating lipotoxicity and promoting aerobic glucose oxidation. Therefore, modulating metabolic reprogramming in cardiomyocytes represents an effective strategy to delay pathological remodeling.

Here, we reveal SF3A2 as a novel deacetylated substrate of SIRT7, uncovering a new mechanism underlying myocardial pathological remodeling. SIRT7 is a nucleolar-enriched, 400-amino acid protein encoded by a 10-exon transcript at chromosome 17q25.3, which is particularly abundant in the heart^33^. Sirt7 deficiency results in a significant reduction in both mean and maximum lifespan in animal models, culminating in the development of myocardial hypertrophy and inflammatory cardiomyopathy^34^. In addition, previous studies have established that SIRT7, an NAD⁺-dependent deacetylase, regulates pathological process by deacetylating diverse substrates. Sirt7 deacetylates Keap1, thereby promoting Nrf2 dissociation and nuclear translocation to activate antioxidant gene expression^35^. Khoa A, Tran et al. reported that SIRT7 degradation via the proteasome pathway promotes NUCKS1 acetylation, enhancing its chromatin binding and thereby inducing metabolic and inflammatory gene expression^36^. Here, we identify SF3A2 deacetylated by SIRT7 conservatively attenuates myocardial pathological remodeling. Under pathological stress, the hearts exhibit reduced NAD⁺ biosynthesis and elevated acetyl-CoA levels, concomitant with diminished SIRT7 activity. The SIRT7 inactivity acetylates SF3A2, leading to specifically skipping of Nnt-Δe2. Consequently, decreased Nnt expression disrupts NAD⁺ regeneration, forming a negative feedback loop that further depletes cellular NAD⁺ pools and exacerbates metabolic reprogram in the cardiomyocytes. Furthermore, our high-throughput screening revealed that ginsenoside Rg1 binds to and activates SIRT7, improving heart function. These findings collectively establish the SIRT7-SF3A2 axis as a key therapeutic pathway for myocardial pathological remodeling.

As a key regulatory mechanism essential for cell fate determination, AS governs over 90% of human genes, enabling cells to generate vast protein diversity from a limited number of genes^37, 38^. Acetylation of splicing factors modulates AS and is implicated in complex diseases, yet whether a conserved regulatory mechanism exists in myocardial pathological remodeling remains an open question. SRSF1, as a splicing factor, regulates the alternative splicing of Bcl2L12, thereby exerting an anti-apoptotic effect via the p53 pathway^39^. RBMS1 promotes cardiac fibrosis by governing the AS of LMO7, which in turn activates the TGF-β1 pathway^40^. SF3A2, implicated in clinical onset and prognosis of MI, modulates the AS of mitochondrial function genes in myocardial I/R injury^41–43^. We found that SF3A2 activity is directly controlled by its acetylation and that deacetylating SF3A2 during the ischemic and I/R phase significantly delayed myocardial pathological remodeling. As a core splicing factor within the U2 snRNP complex, SF3A2 is recruited to pre-mRNA to recognize the branch point sequence and 3’ splice site, enabling the assembly of a catalytic spliceosome^44^. Our findings identify that acetylation of SF3A2 drives its interaction with Ddx1 to promote an exon-skipping event of exon 2 in Nnt. We identify a conserved mechanism in cardiomyocytes whereby SF3A2 deacetylation governs Nnt AS, thereby modulating metabolic reprogramming to attenuate myocardial pathological remodeling.

## Limitations

While uncovering the mechanisms of the SIRT7-SF3A2-Nnt axis in myocardial pathological remodeling, this study has several limitations. First, the reliance on fresh heart tissue for detecting SF3A2 acetylation hinders its validation in clinical patient samples. Future validation experiments with human cardiac organoids generated by differentiating pluripotent stem cells are warranted to confirm SF3A2 acetylation in myocardial pathological remodeling. Second, although Ddx1, which is newly identified here as an SF3A2-interacting protein, is known to function in RNA splicing and protein synthesis, its specific role in pathological remodeling remains unclear. Third, while our study demonstrates the role of SF3A2 in cardiomyocytes, it remains unclear whether it mediates crosstalk with other cell types, such as Tregs and macrophages.

## Conclusions

In summary, we have identified an SF3A2 acetylation-dependent regulatory mechanism in myocardial pathological remodeling. Under pathological remodeling conditions induced by MI and hypertrophy, SF3A2 is acetylated at lysin 10 and promotes the AS of Nnt exon 2, thereby suppressing Nnt expression and NAD^+^ level, and ultimately lead to metabolic reprogramming. AAV9-cTNT mediated SF3A2(K10R) delivery redirects glucose flux toward TCA cycle over anaerobic glycolysis and promote fatty acid β-oxidation to mitigate heart dysfunction. Moreover, we found that SF3A2 is a critical substrate in SIRT7-mediated deacetylation and provide a potential mechanism by which SIRT7 acts as a protective factor in pathological remodeling. In addition, we identified a novel and potent SIRT7 activator that prevents myocardial pathological remodeling, both in cultured cardiomyocytes and in mice, by inhibiting CD36 translocation-mediated lipid toxicity and enhancing OXPHOS function. These observations support the potential utility of targeting SF3A2 deacetylation at K10, or activating SIRT7, as a therapeutic strategy for myocardial pathological remodeling.

### Disclosures

None.

### Supplemental Material

Supplemental Methods

Supplemental Table S1–S3

Supplemental Figures S1–S9

Supplemental References 1–7

